# Bioengineering cobalt chromium cardiovascular stent biomaterial for biofunctionalization

**DOI:** 10.1101/042770

**Authors:** Thamarasee M. Jeewandara

## Abstract

Suboptimal biocompatibility of cardiovascular stents manifest as non-compliance at stent-artery interface *in vivo*. We optimized a plasma-activated coating (PAC) technology to modify cobalt chromium alloy L605 (PAC-L605) surface of an implantable coronary stent material, for improved biofunctionalization. The PAC-L605 surfaces displayed covalent binding capacity of a protein candidate tropoelastin (TE) by retaining 70.3% of TE after SDS detergent washing. Human coronary artery endothelial cell (HCAEC) proliferation visualized with crystal violet staining, did not vary significantly among the biomaterials at 3 or 5 days. Anchorage of cell cytoskeleton visualized with immunofluorescence and scanning electron microscopy (SEM), showed homogenous cell morphology on PAC/TE (with TE) surfaces. Surface hemocompatibility was assessed with static and flow blood assays, the hydrophilic PAC-L605 displayed lower clot formation compared to L605. Area of surface fibrinogen deposited was significantly lower on PAC-L605 vs. L605. Selected ISO 10993-4 tests for biological evaluation of medical devices in contact with blood indicated significantly lowered plasma markers of thrombin-antithrombin complex (TAT), beta-thromboglobulin (β-TG), soluble P-selectin and soluble terminal complement complex (SC5b-9) on PAC-L605 vs. L605. There was no significant difference for plasma biomarkers of polymorphonuclear elastase (PMN elastase) on PAC-L605 vs. L605. Improved surface biofunctionalization of implantable cardiovascular materials could be achieved by plasma-activated coating (PAC).

## Introduction

Evaluating biological design from an engineering perspective, aid researchers develop medical device surfaces with improved biocompatibility for clinical applications. Micro/nanostructured surfaces are developed to improve biomimicry during interaction with micro/nanoscale living tissue (Coutinho et al. 2011). In bioengineering, ‘biofunctionalization’ is the modification of a material to have biological function (Nyanhongo, Steiner, and Gübitz 2011), and elicit appropriate responses during its applications simulated *in vitro* and translated *in vivo*. Key limitations post stent implantation include thrombosis, neointimal hyperplasia (NIH), restenosis and late stent thrombosis (LST) (Cutlip et al. 2007). Cardiovascular stent surface modifications aim to overcome these limitations via optimized interactions with cells, protein and blood at the artery-material interface, reviewed elsewhere (Jeewandara, Wise, and Ng 2014).

The advent of second generation drug-eluting stents (DES), caused significant reduction to the incidence of restenosis and late myocardial infarction (MI) (Alhejily and Ohman 2012). However, among patients with clinical complications, risk of restenosis, stent thrombosis and possible MI still exists (Alhejily and Ohman 2012). Tissue engineering hydrophilic surfaces for research applications aim to; prevent surface aggregation of proteins whilst facilitating controlled protein attachment (Malmsten 1998, Waterhouse et al. 2011, Yin et al. 2009), assist anchorage of healthy cell cytoskeleton on materials for endothelialization (Wozniak et al. 2004), promote surface hemocompatibility (Prentner et al. 2010) and promote biofunctionalization of medical devices in contact with blood in accordance to ISO 10993-4 tests (Seyfert, Biehl, and Schenk 2002).

The present study utilized a proprietary applied plasma physics technique (Bilek et al. 2011) to modify the surface of cardiovascular stent material alloy L605. The plasma-activated coating (PAC) constructs a buffer layer at the interface of organic and inorganic materials forming a robust surface; characterized previously on PAC-L605 (Jeewandara 2016). The surface modification is unique, non-delaminating and able to covalently bind tropoelastin (TE), a bioactive protein candidate of interest (Waterhouse et al. 2011). In the present study, we investigated biofunctionalization of PAC-L605 vs. L605 via 4 aims:

1. Comparative covalent binding capacity of tropoelastin (TE) to surfaces.
2. Human coronary artery endothelial cell (HCAEC) proliferation to determine endothelialization on PAC-L605 vs. PAC-L605/TE (with tropoelastin) vs. L605.
3. Hemocompatibility assays to determine thrombogenicity of PAC-L605 vs. PAC-L605/TE vs. L605.
4. Enzyme linked immunoabsorbent assays (ELISA) to determine concentration of biochemical markers of thrombosis, platelet, complement and leukocyte activation, after blood-material contact (table 1).

**Table 1:** ISO 10993-4 selected tests for biological evaluation of medical device material interaction with blood.

| **ELISA test** | **Molecular marker/cascade investigated** |
| --- | --- |
| Thrombin-Antithrombin complex (TAT) | Thrombosis/Coagulation |
| Beta-Thromboglobulin (β-TG) | Platelet activation |
| Soluble P-selectin (sP-selectin) | Platelet activation |
| Soluble terminal complex C5b-9 (SC5b-9/TCC) | Complement activation (immunology) |
| Polymorphonuclear neutrophil elastase (PMN elastase) | Leukocyte activation (hematology) |

## Materials and Methods

### Preparation of cobalt chromium alloy L605 for surface modification

Samples of alloy L605 (10 cm x 8 cm x 0.025 cm) were prepared as before (Jeewandara 2016). All experiments were carried out with freshly made PAC, used within 2 weeks, unless stated otherwise.

### Synthesis of plasma-activated coating

The cleaned substrates were mounted onto a stainless steel cylindrical ccrf reactor. Base pressure of the system was pumped down to 10^−6^ - 10^−5^ Torr. A reactive mixture of argon, nitrogen and acetylene plasma was introduced in the upper part of the reactor. The flow, Q, of each gas for alloy L605 surface; Q_Argon_=3 sccm, Q_Acetylene_=1 sccm, and Q_Nitrogen_=10 sccm (500 V, t=20 min), was controlled individually, using mass flow controllers and maintained constant, during coating deposition (PAC1 recipe). On another alloy L605 sample of similar dimensions, we trialed a second recipe Q_Argon_ = 3 sccm, Q_Acetylene_ = 2 sccm and Q_Nitrogen_ = 10 sccm (1000 V, t = 10 mins) (PAC2 recipe). Plasma was sustained at a total pressure of 80 mTorr, rf power 50 W, and deposition time of 10-20 mins. Unless specified, two recipes of PAC (PAC1/PAC2) were investigated simultaneously.

### Tropoelastin (TE) coating and SDS-ELISA detection

Recombinant human tropoelastin (TE) used in previous studies (Waterhouse et al. 2010) was similarly expressed and purified from E.coli expression system (Martin, Vrhovski, and Weiss 1995). The PAC-L605 and alloy L605 samples (0.8 cm x 0.8 cm) were incubated with 250 μl of 50 μg/ml tropoelastin in phosphate buffered saline (PBS), at 4°C overnight. SDS detergent treatment and ELISA for characterization of bound TE was carried out as previously described (Yin et al. 2009).

### Cell culture and proliferation

Cell assays were performed in triplicate in tissue culture plastic (TCP) 24-well plates containing samples of L605, PAC-L605 and PAC-L605/TE (0.6 cm x 0.6 cm). HCAECs were seeded and left to proliferate for 3 or 5 days. Cell proliferation was quantified by crystal violet staining and measurements of absorbance at 570 nm.

### Immunofluroscent detection of cell cytoskeleton

Metallic samples with or without tropoelastin (50 μg/ml) were seeded with HCAECs. Five biomaterial surfaces were investigated: 1. PAC-L6051 recipe, 2. PAC-L6052 recipe, 3. PAC 1 recipe with tropoelastin (TE), 4. PAC 2 recipe with TE, 5. Cobalt chromium bare metal alloy L605. After day 5 cell culture, samples were incubated with 0.2 M glycine for 20 min and fixed in triton X-100 (0.2% v/v in PBS) for 10 min. Cells were blocked in 5% (w/v) BSA in PBS for 1 hour. Samples were stained with human anti-vWF antibody (Dako, 1:400 in PBS) for 30 min, washed thrice in PBS and incubated with goat anti-rabbit IgG Alexa Fluor^®^488 (Abcam, 1:90 in PBS) for 1 hour. Samples were stained with phalloidin-TRITC (Sigma, 0.1 μg/ml in 3% BSA) for 45 min washed thrice with PBS and stained with DAPI (Sigma, 0.15 μg/ml in PBS) for 15 min. Samples were imaged with an Olympus IX70 inverted microscope using a 40x objective.

### Scanning Electron Microscopy (SEM) detection of cell cytoskeleton

HCAECs were cultured for 2 days on three sample surfaces: L605, PAC-L605 and PAC-L605/TE. Samples were washed in PBS, fixed in 4% formaldehyde for 30 min at room temperature (RT), dehydrated in ascending grades of ethanol (30% v/v to 100% v/v, 5 min each) before chemical drying with hexamethyldisilazine (HMDS). The samples were mounted to stubs using carbon tape, sputter coated with 20 nm gold and imaged with Zeiss Ultra field emission scanning electron microscope (FESEM).

## Hemocompatibility assays

### Blood sampling and platelet rich plasma (PRP) isolation for static assay

Whole blood was obtained as previously described (Waterhouse et al. 2010). All experiments were conducted in triplicate samples at least thrice with different donor’s blood. Samples of L605, PAC-L605 and PAC-L605/TE (0.8 cm x 0.8 cm) were placed in a 24 well plate blocked with 3% BSA in PBS for 1 hour at RT and washed thrice with PBS. Blood was anti-coagulated with 0.5 U/ml heparin and centrifuged for 15 min at 1000 rpm. The supernatant was removed and samples incubated with 250-500 μl of PRP for 60 min at 37°C while rocking. Samples processed for SEM as above were additionally post-fixed with OsO4 (ProSciTech, 0.1% in 0.1 M PB) for 1 hour, prior to dehydration process.

### Ageing study on thrombogenicity

An ageing study was conducted with PRP to analyze surface thrombogenicity over time. The samples were 1) 1-month old PAC-L605, 2) 1 month old PAC-L605 with TE stored for 2 weeks at −20°C, 3) 1 month old PAC-L605 incubated with TE stored overnight at 4°C and 4) bare alloy L605 (control). Samples were placed in a 24 well plate and incubated with PRP at 37°C for 60 min, and processed for SEM imaging.

### Whole blood adhesion assays

Samples PAC-L605 and L605 (0.8 cm x 0.8 cm) were incubated with 500 μl of heparinized whole blood (0.5 U/ml) for a period of 30, 60, and 90 min at 37°C while rocking, to assess variation of surface hemocompatibility with time of flow. Samples were processed for SEM imaging.

### Whole blood flow assays – Modified chandler loop system

Coagulation under flow was investigated with a modified chandler loop system at physiological conditions *in vitro* as previously described (Gaamangwe 2014, Waterhouse et al. 2010). The blood sample volume and dimensions of all material samples met ISO 10993-12 regulations (Gaamangwe 2014). Clot formation was quantified for sample surfaces of; 1-week old PAC recipe 1 vs. PAC recipe 2 vs. L605, followed by PAC-L605 vs. PAC/TE vs. L605, and 5-month old PAC-L605 vs. L605. Plasma from whole blood was separately isolated for surface fibrinogen detection and ISO-10993-4 selected ELISA tests.

### Surface Fibrinogen detection

Fibrinogen staining helped visualize surface fibrin deposited on biomaterials after blood flow assay. A stock solution of fibrinogen Alexa Fluor^®^488 was prepared (Life Technologies, 5 mg fibrinogen in 3.3 ml of 0.1 M NaHCO3, pH 8.3, at RT) and gently agitated for 1 hour. Samples (PAC-L605 and L605) were removed from chandler loop and incubated in a 24-well plate with a volume of 250 μl fibrinogen-Alexa Fluor^®^488 (1:60 dilution in distilled water) for 15 min. The samples were washed with PBS and incubated with DAPI (Sigma, 0.15 μg/ml in PBS) for 15 min. Samples were imaged with an Olympus IX70 inverted microscope using a 10x objective. Surface fibrinogen on 5 areas per sample image, were quantified and averaged with ‘smart segmentation’ tool, Image Pro Premier software (Media Cybernetics, USA).

### ELISA for biomarkers of coagulation, inflammation and complement activation

Whole blood samples from chandler loop assay were centrifuged at following conditions to isolate plasma prior to ELISA tests (table 2).

**Table 2:** Centrifuge conditions of plasma isolation for individual ELISAs.

| ELISA test for biomarker | Plasma Isolation Conditions |  |  |
| --- | --- | --- | --- |
|  | Centrifugation Speed (x g) | Time (min) | Anticoagulant |
| Thrombin Antithrombin (TAT) | 3000 | 10 | 10% citrate |
| Beta thromboglobulin (β-TG) | 1000 | 20 | 10% citrate |
| Soluble P-selectin | 1000 | 15 | 10% citrate |
| SC5b-9 complex | 10,000 | 5 | 10% K<sub>2</sub>EDTA |
| PMN elastase | 1500 | 10 | 10% K<sub>2</sub>EDTA |

The concentrations of TAT complex (abcam, Cambridge, MA, USA), β-TG (MyBioSource, San Diego, CA, USA), sP-selectin (LifeTechnologies, Vic, Aus), SC5b-9 terminal complement complex (MicroVue, Quidel, San Diego, CA, USA) and PMN elastase (abcam, Cambridge, MA, USA) were quantified according to manufacturer’s protocol. Plasma biomarkers of sP-selectin, SC5b-9 and PMN elastase were respectively diluted by factors of 1:30, 1:10 and 1:100 in specimen diluent. Manufacturer-provided high and low controls for SC5b-9 and PMN elastase, were used in ELISA tests in similar dilution to the samples. Concentration readings to identify presence of biomarkers - TAT was performed at 450 nm with a 570 nm wavelength correction, β-TG at 450 nm, sP-selectin at 450 nm with a 620 nm wavelength correction, TCC/SC5b-9 complex at 450 nm and finally PMN elastase at 450 nm with a 620 nm wavelength correction. All experiments were conducted in triplicate samples.

## Statistics

Data are expressed as mean ± standard deviation. Data were analyzed by paired t-test and ordinary one-way ANOVA with multiple comparisons for >2 variables. For ELISA test results, the unknown concentrations were calculated with nonlinear regression (curve fit). Standard curves to interpolate hyperbola (X is concentration) were selected with fitting method (least squares ordinary fit) to interpolate unknowns, using GraphPad Prism version 6.00 (Graphpad software, San Diego, CA, USA) for MAC OS X.

## Results

Covalent protein binding capacity of PAC surfaces were determined using an elastin-specific ELISA. In the absence of tropoelastin (TE), background levels of absorbance were 0.012±0.009 on PAC-L605 and 0.010±0.014 on alloy L605 respectively (Figure 1). Absorbance readings without SDS washing for PAC-L605 vs. L605 was 0.202±0.01 and 0.138±0.005 respectively. SDS washing was used to remove non-covalently bound protein, after which readings decreased to 0.142±0.011 for PAC-L605 (29.70 % reduction) and 0.041±0.009 for L605 (70.28% reduction). The modified PAC-L605 displayed covalent binding capacity by retaining 70.3% of bound TE after SDS wash.

**Figure 1:**
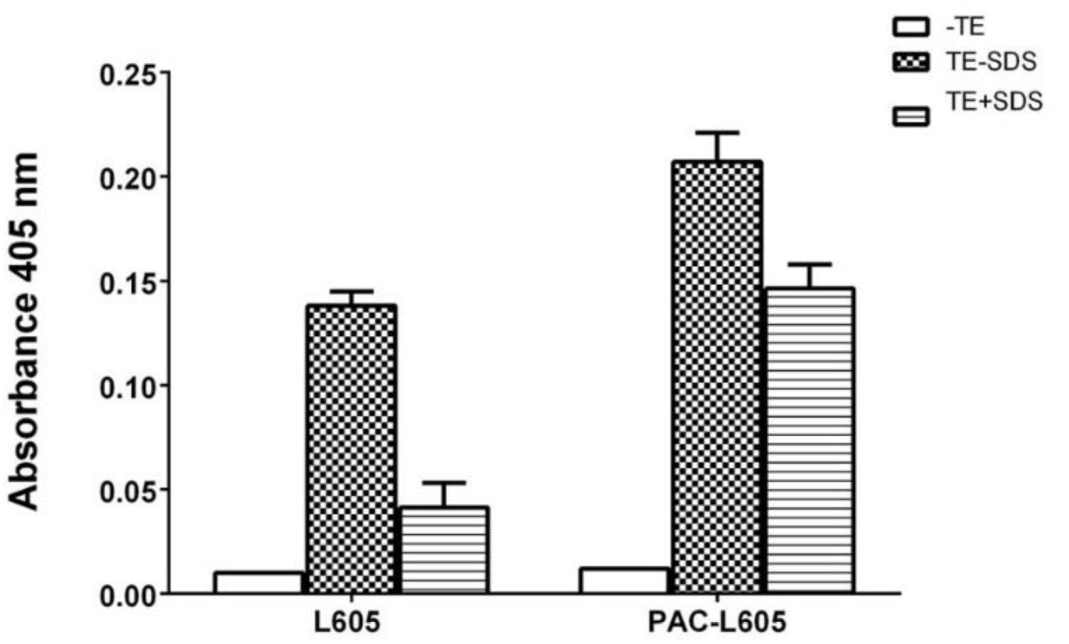
ELISA with anti-elastin antibody to detect covalent binding capacity of TE to plasma-modified PAC-L605 surfaces, compared to bare metal alloy L605. The TE-SDS samples (checked) were not washed with SDS detergent, the TE+SDS samples (stripes) were washed with SDS detergent. The blanks are negative controls.

### Proliferation and anchorage of HCAECs: cell-material interactions

HCAEC proliferation at day 3 and day 5 were quantified on biomaterials L605, PAC-L605 and PAC-L605/TE (with TE). Tissue culture plastic (TCP) was the positive control. At day 3, absorbance readings at 570 nm for TCP vs. L605 vs. PAC-L605 vs. PAC-L605/TE were 0.272±0.034, 0.126±0.012, 0.135±0.013 and 0.119±0.009 respectively. At day 5, absorbance for material surfaces in the same order were 0.535±0.017, 0.194±0.014, 0.271±0.016 and 0.304±0.048 respectively. Cell proliferation was significantly higher for TCP (positive control, not shown), although not significantly different among biomaterials of interest at day 3 or 5 cell culture (Figure 2).

**Figure 2:**
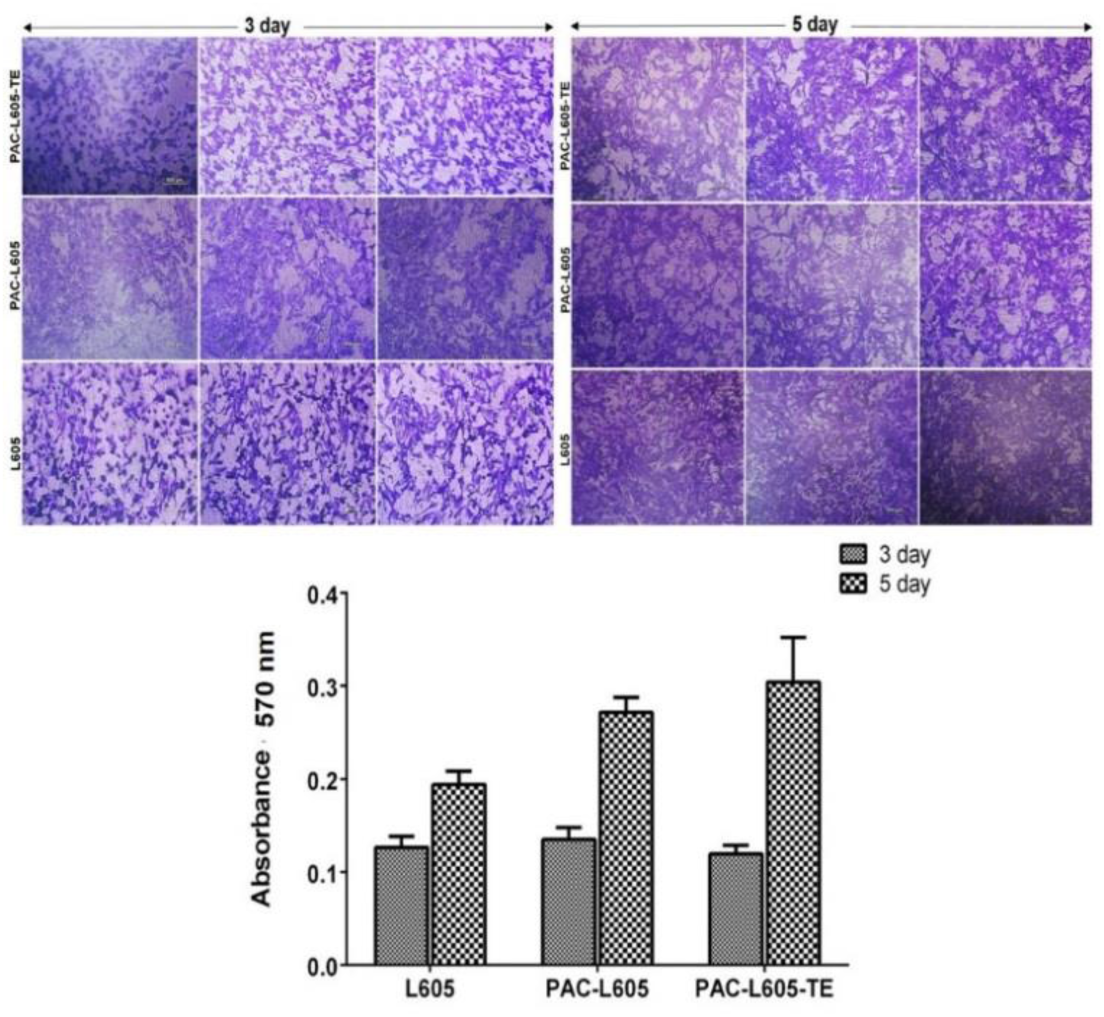
Figure panels on top indicate crystal violet stained HCAEC proliferation on PAC-L605/TE, PAC-L605 and alloy L605, at day 3 and day 5 respectively (10 x magnification, scale bar 200 μm). The graphed absorbance of surface cell density increased progressively from day 3 to day 5, although results were not significantly different among the biomaterials at either time point of study.

Filamentous actin (F-actin) anchorage of HCAECs to biomaterials were visualized with inverted fluorescence microscopy on 5 different biomaterial surfaces at day 5 proliferation (40 x magnification). Homogenous distribution of F-actin was observed on PAC2-L605 surfaces compared to PAC1-L605. Similar trends were also observed with PAC2-L605 with TE (Figure 3). The PAC2 recipe was chosen for further cell experiments hitherto referred as PAC-L605.

**Figure 3:**
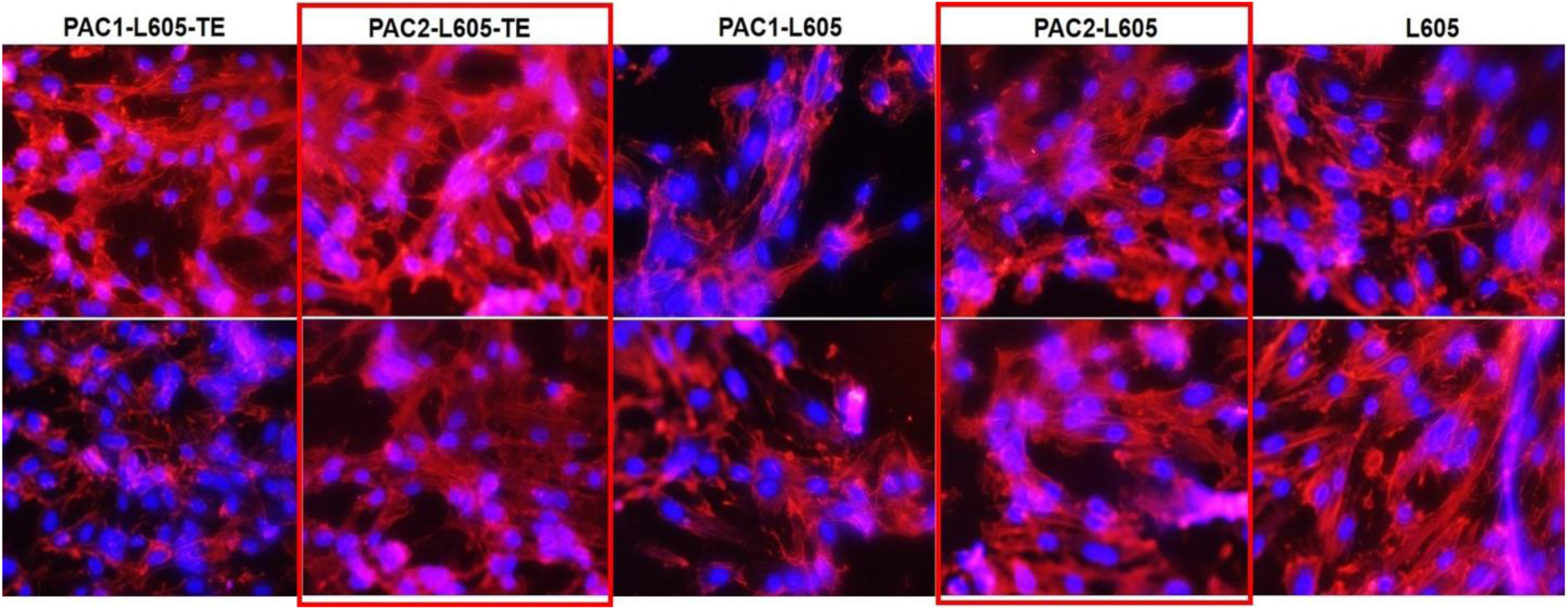
The F-actin anchorage on biomaterials were visualized with phalloidin-TRITC staining at 40 x magnification (fluorescent channel, inverted Olympus microscope). The cytoskeleton spread homogenously (F-actin) on PAC2 surfaces, with or without tropoelastin (TE), recipe chosen for further experiments in cell culture.

Distribution of F-actin, vWF and cell nuclei (triple stain) were visualized for HCAECs on PAC-L605 vs. L605 (40 x magnification). Endothelial cells on PAC-L605 appeared more confluent and homogenous in distribution compared to HCAECs on L605.

**Figure 4:**
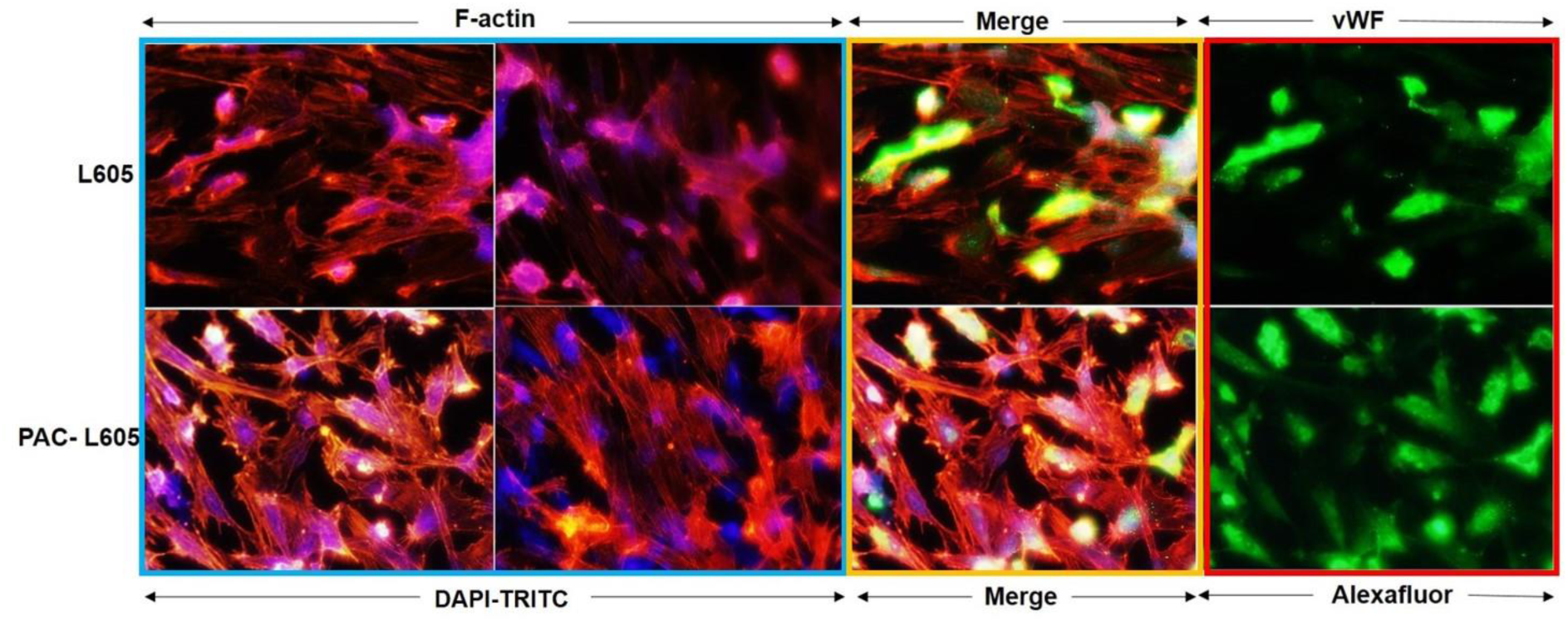
Filamentous actin visualized on PAC-L605 and L605 surfaces at 40x magnification with phalloidin-TRITC and DAPI after day 5 proliferation. Image shows well defined cytoskeleton on PAC-L605 compared to L605 (blue box). Merged triple stain (yellow) and endothelial specific vWF staining uptake (red box).

The HCAEC morphology on L605, PAC-L605 and PAC-L605/TE material surfaces were visualized after day 2 proliferation with scanning electron microscopy (Figure 5). The PAC-L605/TE (with tropoelastin) showed least spindle projections, homogenous cell cytoskeleton distribution and support for HCAEC proliferation.

**Figure 5:**
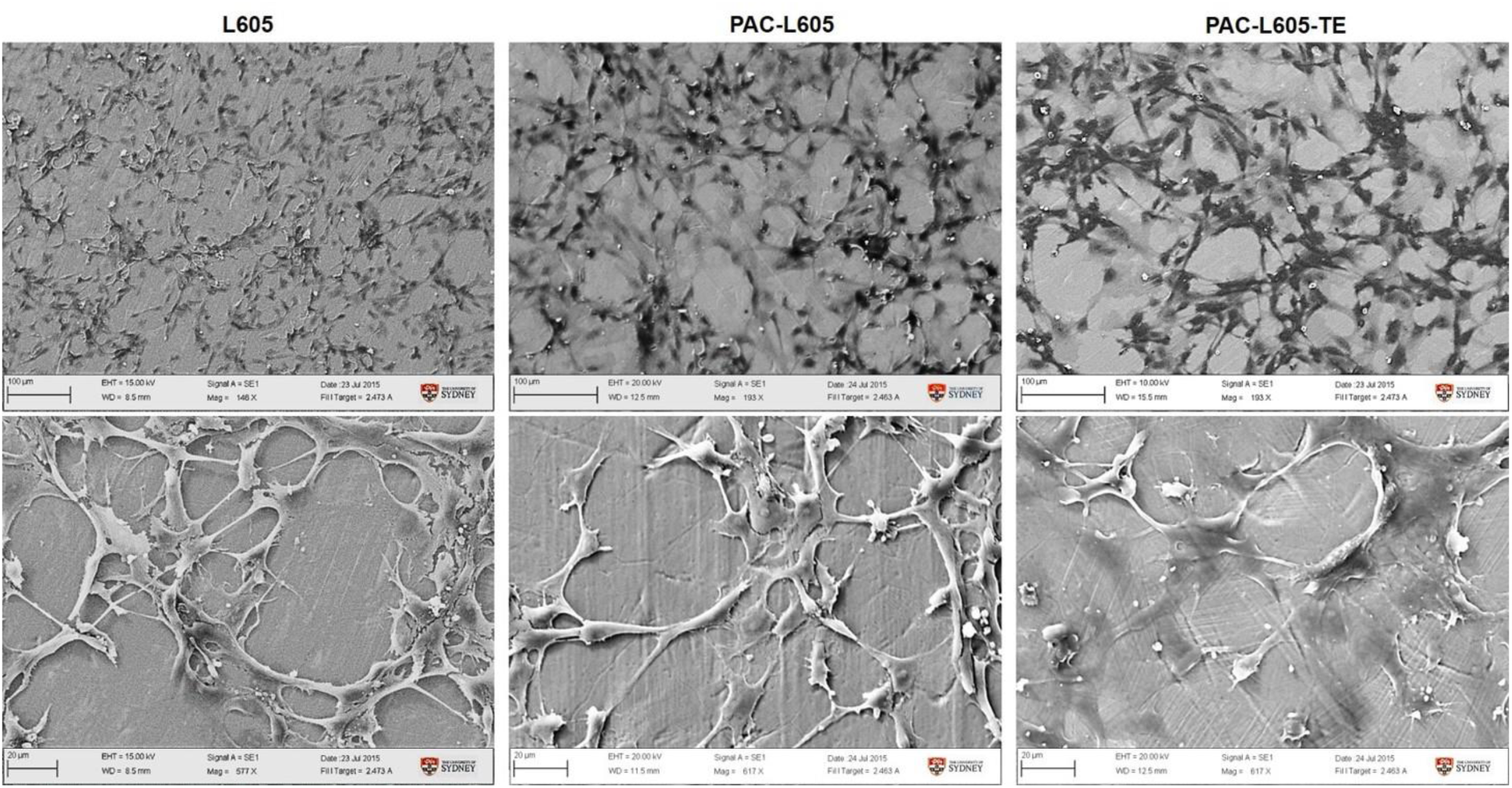
HCAECs on alloy L605 and PAC-L605 show more spindle projections compared to PAC-L605/TE surfaces. The PAC+TE surfaces supported confluent, homogenous HCAEC growth and improved surface-cytoskeleton attachment.

### Hemocompatibility of plasma-modified biomaterials

The presence of protein in PRP, prevented platelet and fibrin adherence to modified, hydrophilic PAC-L605 and PAC-L605/TE surfaces. Fibrins and platelets adhered on the hydrophobic, bare metal alloy L605 (Figure 6).

**Figure 6:**
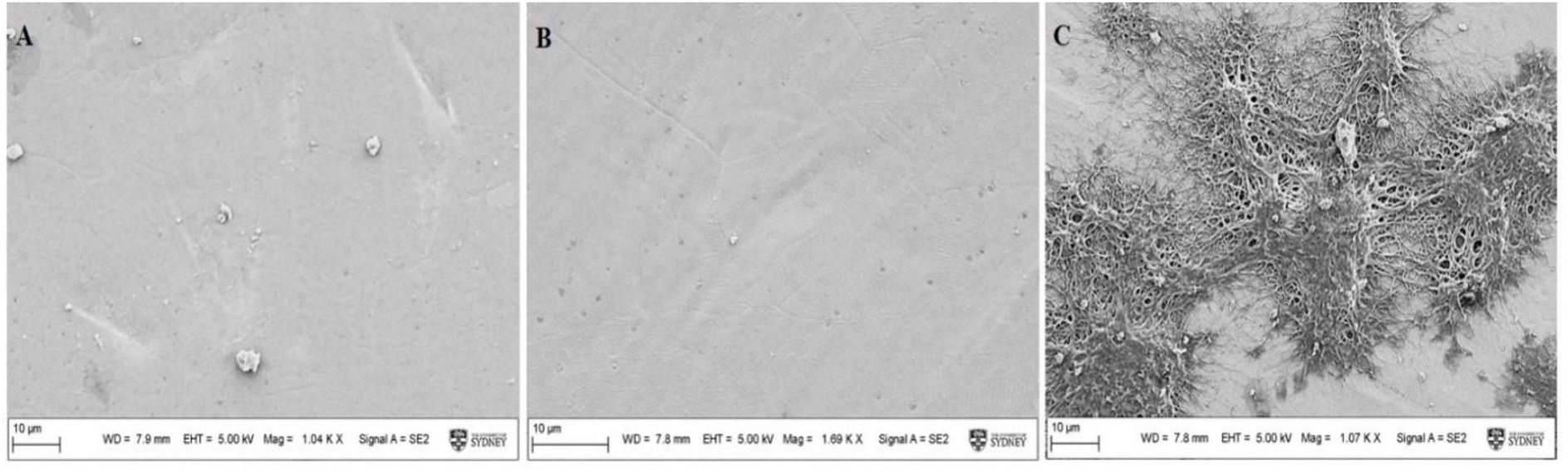
Hydrophilic surfaces PAC-L605 and PAC-L605/TE (A&B) prevented protein aggregation compared to hydrophobic L605 (C), FESEM, 1.2 Kx magnification.

The surface ageing study conducted with PRP showed sustained hemocompatibility for both PAC-L605/TE surfaces incubated with TE for 2 weeks (-20°C) and overnight (4 °C). Hemocompatibility was retained on PAC-L605 surfaces stored air tight for 1 month. Surface platelet aggregation and fibrin clot formation was seen on alloy L605 (Figure 7 A-D).

**Figure 7:**
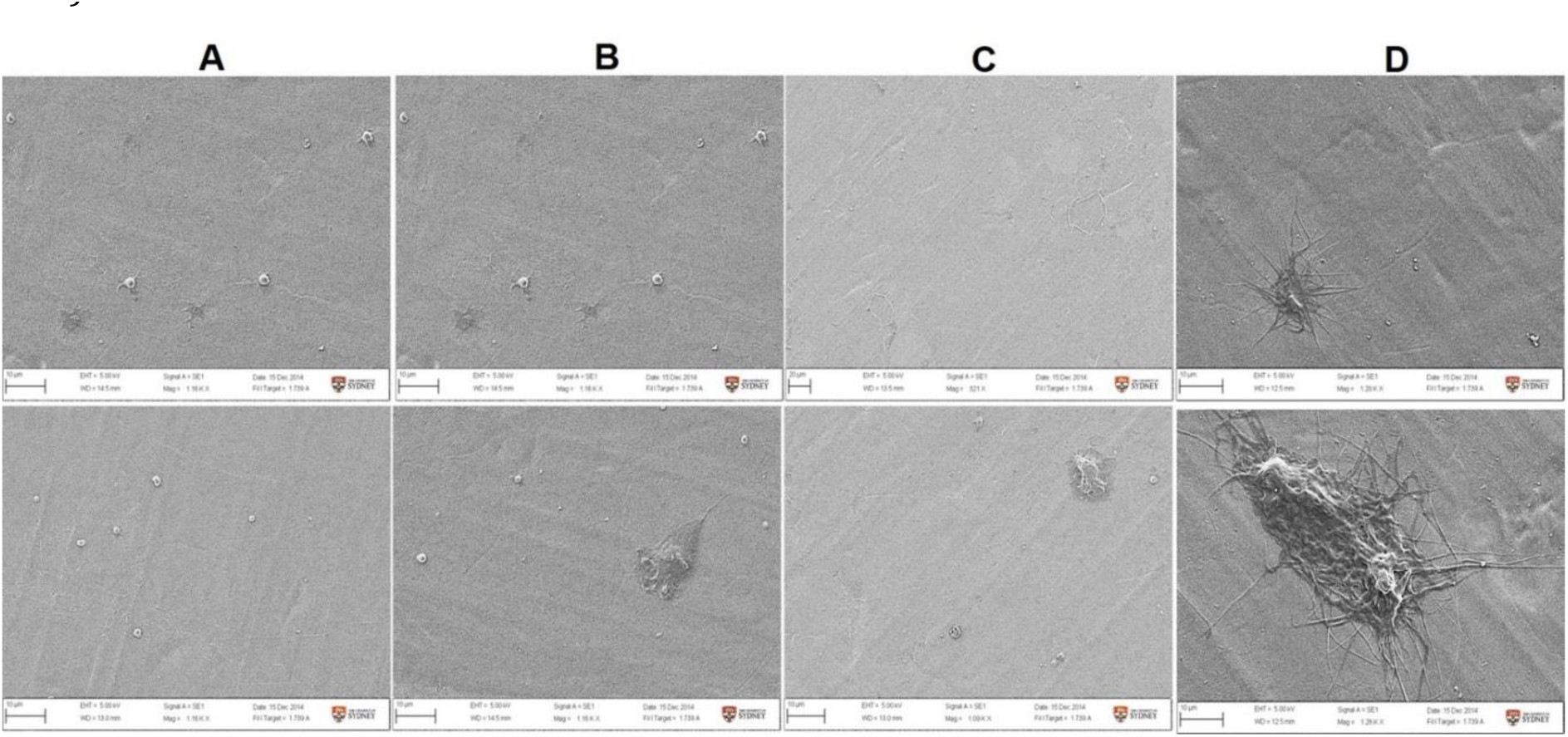
SEM images of the ageing study showed sustained hemocompatibility on A) PAC-L605/TE (2 weeks, −20°C), B) PAC-L605/TE (overnight, 4°C), C) PAC-L605 (air tight, 1 month) and surface platelet/fibrin clot deposition on D) alloy L605. (1.08 Kx magnification)

Whole blood adhesion assays showed lower thrombogenicity for PAC-L605 vs. L605 within the time frame of 30, 60 and 90 min flow. For optimum surface hemocompatibility assessment, the 60 min time-point was selected (Figure 8).

**Figure 8:**
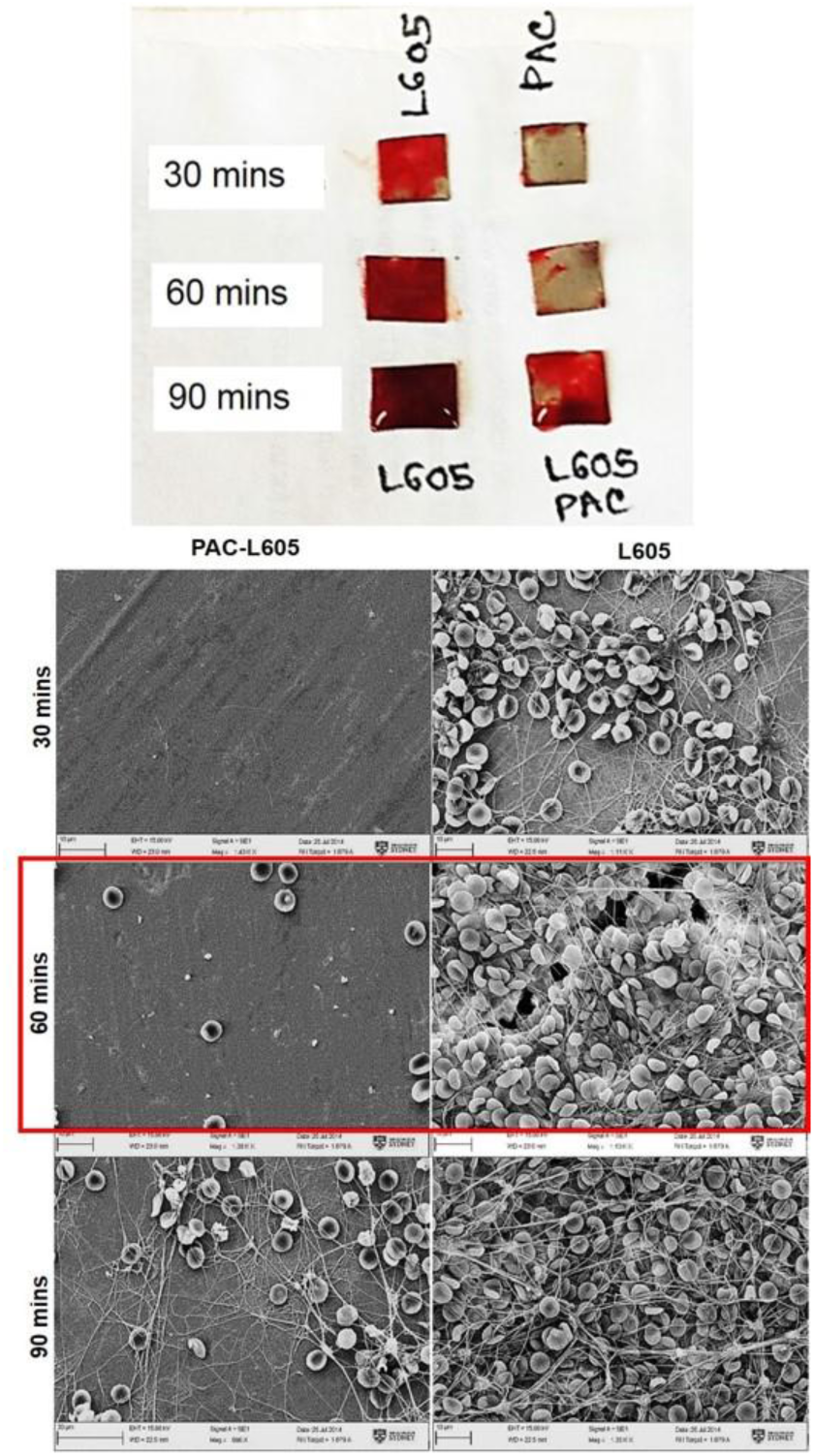
Comparatively lower thrombogenicity observed at 30, 60, and 90 min time points for PAC-L605 vs. L605.

Hemocompatibility under flow conditions with a modified chandler loop demonstrated lowered clot formation for PAC1-L605 vs. PAC2-L605 vs. L605 (2.5±1.1 mg vs. 8.63±2.5 mg vs. 22.2±11.4 mg, p=0.06, not significant) (Figure 9A).

**Figure 9:**
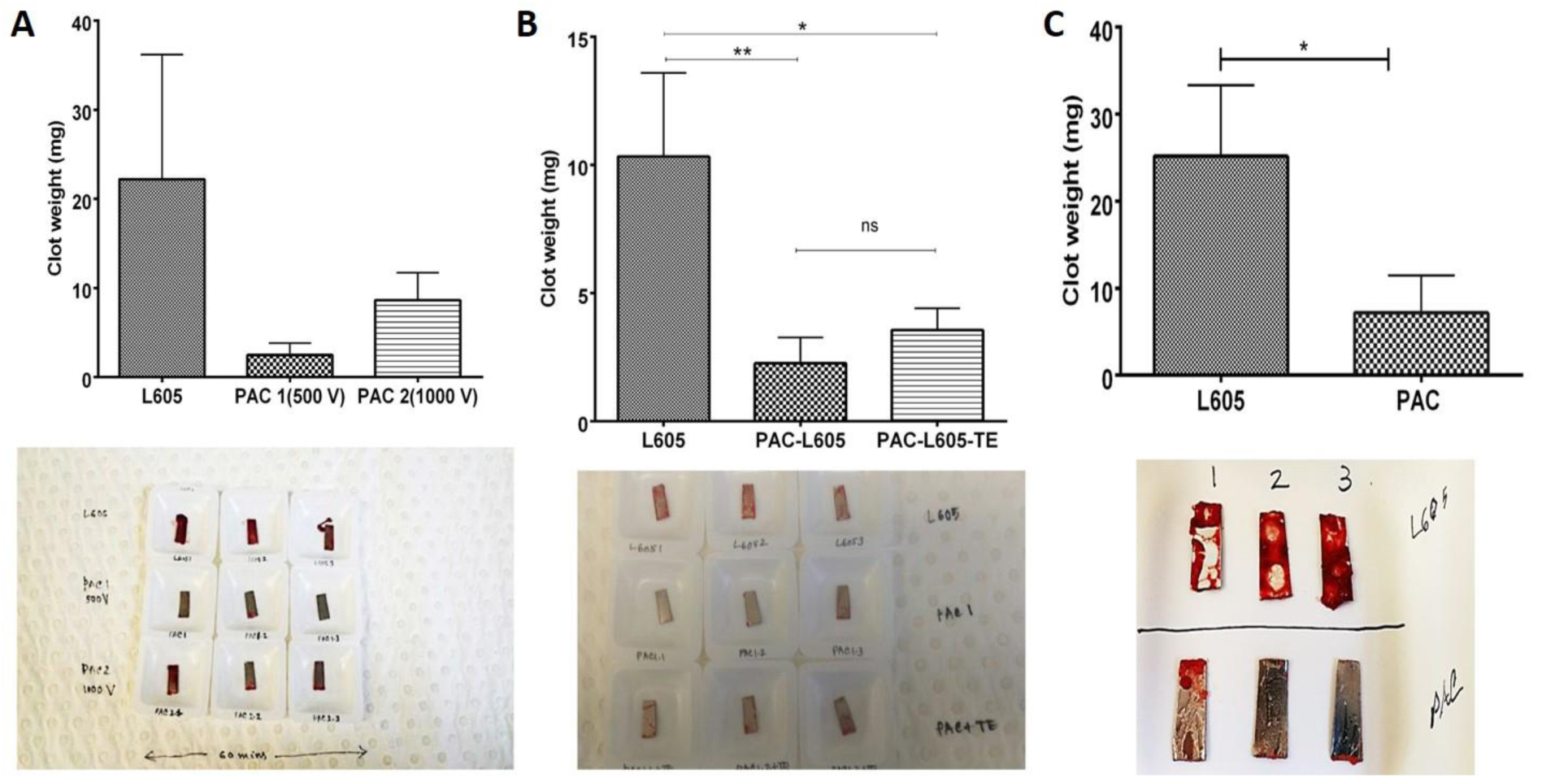
Hemocompatibility under flow conditions A) thrombogenicity among PAC recipes 1&2 compared to L605 B) clot formation reduced for plasma-modified surfaces with/without TE compared to L605 C) surface hemocompatibility retained 5 months after storage for PAC-L605 vs. L605.

Flow assays were repeated with PAC1-L605 recipe (hitherto referred as PAC-L605 for blood assays) vs. PAC-L605/TE vs. L605, 1 week post modification (2.26±1.0 mg vs. 3.56±3.6 mg vs. 10.33±3.3 mg, p<0.05).The presence of tropoelastin did not contribute to significantly lowered clot formation compared to PAC surfaces alone (Figure 9B)

Five months post-modification flow assays were repeated with PAC-L605 vs. L605 (7.16±3.5 mg vs. 25.16±6.66 mg, p<0.05). As with static blood assays, hemocompatibility was retained with ageing (Figure 9C).

Surface fibrinogen visualized after 60 min chandler loop flow assays, indicated lower fibrinogen staining uptake area on PAC-L605 vs. L605 (481.40±263.92 μm^2^ vs. 1451.82±867.83 μm^2^). Increased standard deviation of surface fibrinogen area indicated inconsistent deposition (Figure 10).

**Figure 10:**
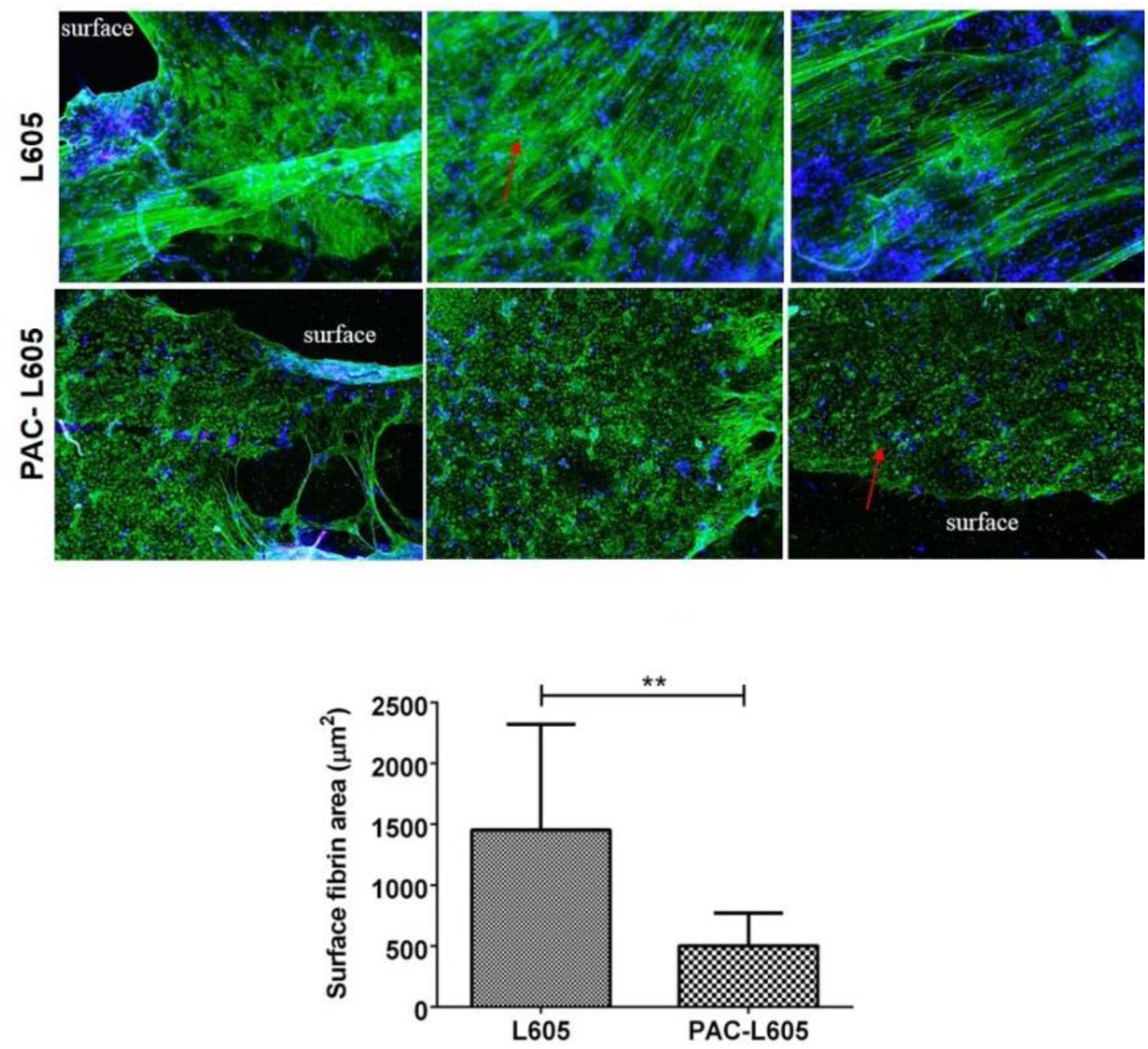
Surface fibrinogen deposition (red arrows) visualized with fibrinogen-Alexa Flour 488 and DAPI (nuclei), on L605 vs. PAC-L605. (10x magnification, 200 μm scale bar). Area calculated with Image Pro Premier software (Media Cybernetics, USA).

### Plasma biomarker activation after blood-material contact

Plasma isolated after 60 min blood-material contact for ELISA, showed significantly lowered biomarkers of TAT for PAC-L605 vs. L605 (7.22±1.33 ng/ml vs. 15.93±2.55 ng/ml, p=0.002) (Figure 11 A). Similar results were observed for β-TG (16.61±3.33 ng/ml vs. 27.16±3.75 ng/ml, p=0.025) (Figure 11B), soluble-P selectin (4.87±0.07 ng/ml vs. 10.575±0.20 ng/ml, p=0.013) (Figure 11C) and SC5b-9 (98.18±21.88 ng/ml vs. 120.27±15.05 ng/ml, p=0.032, high control 113.66±1.41 ng/ml and low control 34.20±1.98 ng/ml) (Figure 11D). Biomarker PMN-elastase was not significantly different for PAC-L605 vs. L605 (3.26±0.26 ng/ml vs. 2.72±0.45 ng/ml, p=0.149, high control 10.67±2.9 ng/ml and low control 1.19±0.41 ng/ml) (Figure 11E). For consistency, all biomarker concentrations were graphed in their original values (Figure 11).

**Figure 11:**
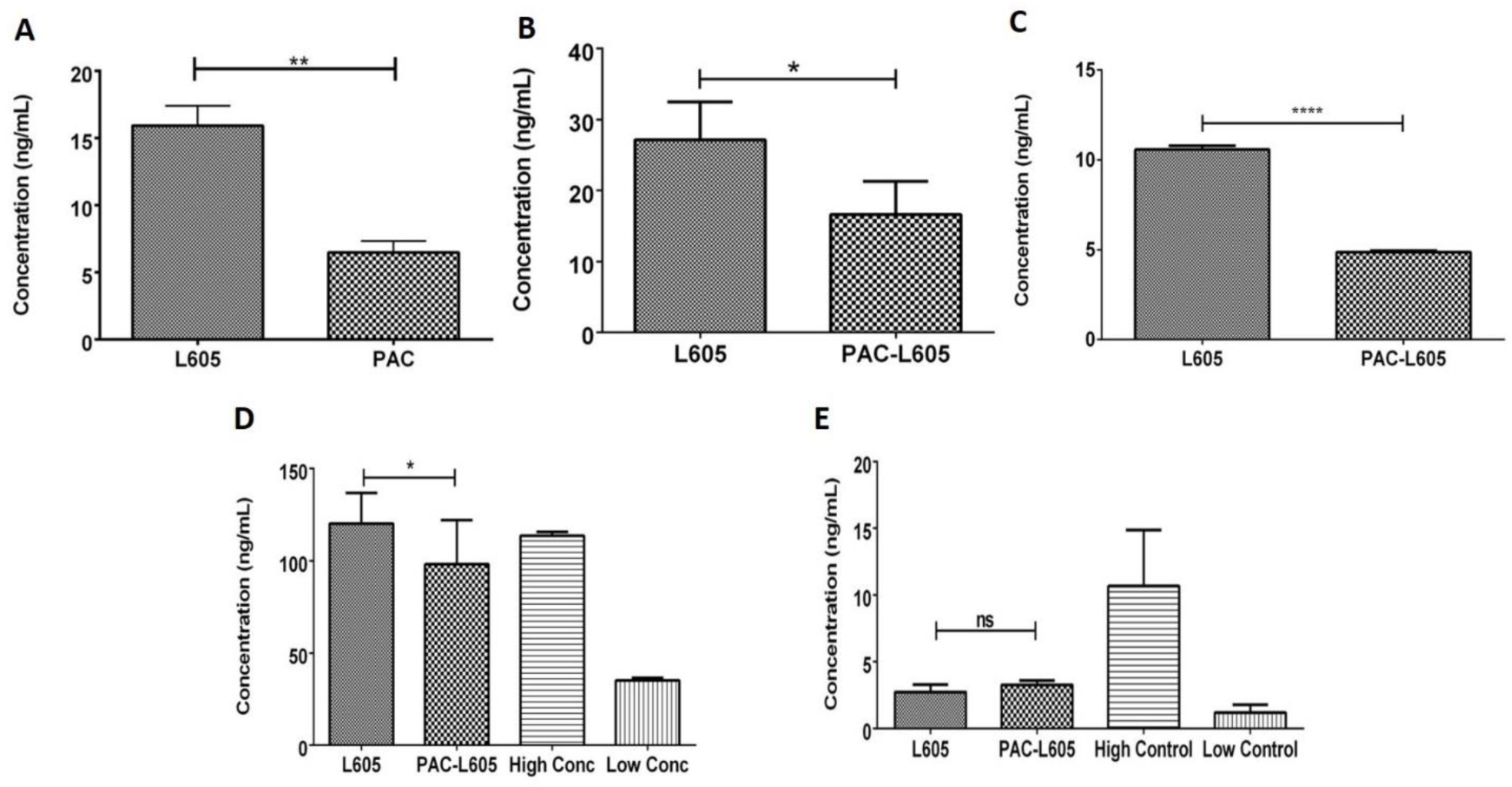
ELISA quantification of biomarkers after blood-material contact for PAC-L605 vs. L605 A) TAT (no dilution, p=0.002), B) β-TG (no dilution, p=0.025), C) sP-selectin (1:30 dilution, p=0.013), D) SC5b-9 (1:10 dilution, p=0.032) and E) PMN elastase (1:100 dilution, p=0.149)

Final concentrations for PAC-L605 vs. L605 after x dilution factor, are noted within parentheses for biomarkers sP-selectin (1:30 dilution; 146.19±2.1 ng/mL vs. 317.25±6 ng/mL), SC5b-9 (1:10 dilution; 981.91±218.84 ng/ml vs. 1202.77±150.53 ng/ml, high control 1136.6±14.16 ng/ml and low control 342.0±19.80 ng/ml) and PMN elastase (1:100 dilution; 272.75±45.17 ng/ml vs. 326.84±26.03 ng/ml, high control 1067.52±296.83 ng/ml and low control 119.66±40.82 ng/ml).

## Discussion

The salient findings of this study, for the previously characterized, plasma-modified PAC-L605 surface are:

1. Plasma-based surface modification allows covalent binding of tropoelastin (TE) in its native conformation.
2. Surface assisted endothelialization at cell-material interface, promotes healthy anchorage of cell cytoskeleton to material for homogenous proliferation.
3. Improved hemocompatibility. Reduced clot formation, fibrinogen deposition and reduced plasma biomarkers of coagulation, complement and platelet activity.
4. Surface biofunctionalization retained with age/time.

The data demonstrates biofunctionalization of plasma-modified surfaces *in vitro*, for vascular compatibility at the artery-stent material interface. The unique, controlled, covalent protein binding capacity on PAC was due to the presence of surface free radicals (Bilek et al. 2011, Waterhouse et al. 2010). However, covalently bound protein surfaces PAC-L605/TE, did not significantly improve endothelialization or hemocompatibility compared to PAC-L605 surfaces alone.

A common technique for clinical engineering biomimetic microsystems is isotropic material surface development with nanoscale/micron roughness to facilitate cell anchorage and growth (Huh et al. 2013, Place, Evans, and Stevens 2009). The cell cytoskeleton is theorized to play an integral role for coordinated attachment to a substrate (Ingber 1993). The SEM and fluorescent images obtained were limited, as they were a 2-dimensional representation of 3-D cell-material organization. However, results indicated assisted, anchorage dependent, cell spreading and proliferation on PAC2-L605 surfaces, with or without TE. Tissue engineering surfaces that mimic physiological environment for biomimicry are one of the most basic principles of biological design.

The *in vitro* model to assess hemocompatibility is an over-simplification of a very complex physiological system (Gaamangwe et al. 2015). The modified chandler loop system was based on previous experimental standardizations (Gaamangwe et al. 2015, Gardner 1974). Practically, hemodynamics play a quantitative role between stent implantation and thrombosis *in vivo* (Foin et al. 2014), and should be taken into consideration during simulations *in vitro*.

Under physiological conditions of static and flow blood assays, platelets aggregate together by thin fibrin strands to form thrombi on hydrophobic L605 in contrast to hydrophilic PAC-L605 surfaces. Similar results were observed with amphiphilic fibrinogen aggregates on L605 compared to PAC-L605. Hemocompatibility of modified surfaces (with or without tropoelastin), during flow and static blood assays, were retained after oxidation/ageing.

The concentration of biomarkers quantified, depend on protein-material interaction during exposure to whole blood. Predominant proteins within extracellular matrix (ECM); albumin, immunoglobulin (IgG), fibrinogen, factor XII and high molecular weight kininogen (HMWK) interact with material surface in a phenomenon termed ‘Vroman effect’ (Wilson et al. 2005). Accordingly, hydrophilic surfaces displace proteins to create a high surface protein turnover rate at material interface, whereas proteins on hydrophobic surfaces do not displace. Instead, surface proteins undergo conformational changes on hydrophobic surfaces, form fibrinogen aggregates, and trigger activation of biomarkers (Vroman and Adams 1969). Biochemical events occurring after blood-material contact to trigger activation of biomarkers, quantified herein, are summarized (Figure 12).

**Figure 12:**
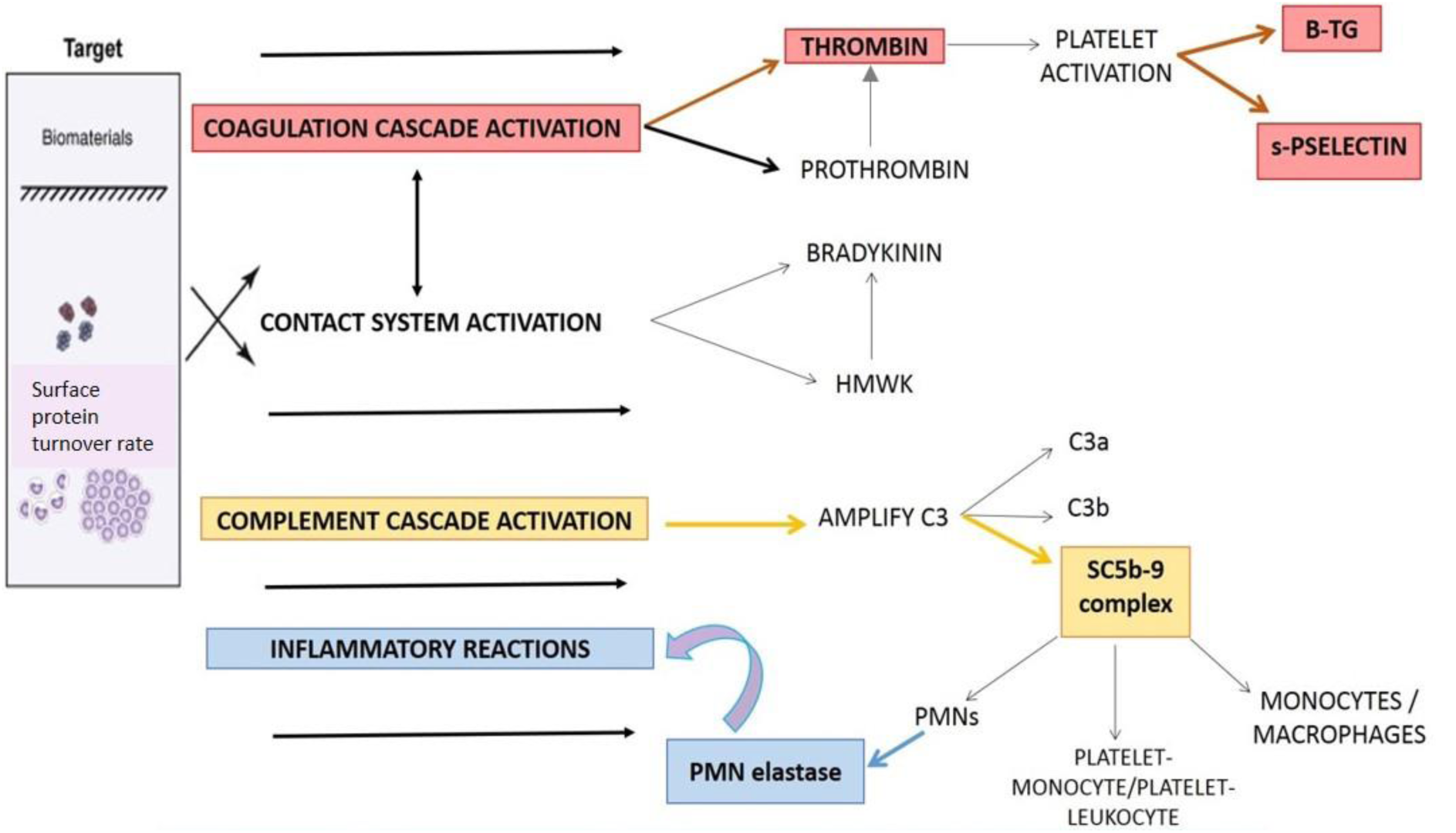
A simple schematic outlining the biochemical pathways activated during blood-biomaterial contact. The highlighted biomarkers were quantified with ELISA tests in this study.

## Conclusion

We investigated biofunctionalization of a previously characterized, plasma-modified, cobalt chromium alloy PAC-L605. Initial studies showed optimum endothelialization for PAC recipe 2 and optimum hemocompatibility for PAC recipe 1. All PAC surfaces covalently bound tropoelastin (TE) compared to L605, but this did not markedly improve biofunctionalization of PAC. With age, biofunctionalization of PAC surfaces retained. For further investigations we propose an optimized recipe of PAC (C-1, N-10, t=10 min, 500 V), which may simultaneously promote both endothelialization and hemocompatibility. Tests in accordance to ISO 10993-4, for biological evaluation of medical devices using PAC recipe 1 vs. L605, indicated comparatively superior hemocompatibility for PAC. The preliminary biofunctionalization study, optimized plasma-activated coating (PAC) on a cobalt chromium material and optimized ISO-109934 recommended *in vitro* biological tests, prior to PAC deposition on a cobalt chromium stent platform for further research.

## Acknowledgements

Manuscript originally produced as thesis chapters (*‘Plasma based biofunctionalization of cobalt chromium alloy L605’* and *‘Biological evaluation of medical devices; selection of tests for interactions with blood’*) for Doctor of Philosophy (PhD) Medicine, Sydney Medical School, University of Sydney – September 2015. Initial research work presented at the Atherosclerosis, Thrombosis and Vascular Biology (ATVB) Scientific conference, American Heart Association, San Francisco, CA, USA, May 2015 (DOI: <u>10.7490/f1000research.1111024.1</u>). The author acknowledges –

*Applied Plasma Physics Department of the University of Sydney, for plasma-activated coating (PAC) development*.

*The Australian Center for Microscopy and Microanalysis (ACMM), University of Sydney, for research and technical support; Mr. Steven Moody (SEM Specialist), Dr. Patrick Trimby (SEM Specialist) and Miss Naveena Gokoolparsadh (Biological Specimen Preparation Specialist)*

*The Heart Research Institute, Newtown, Tissue Culture lab facilities, research and technical support; Pat Pisansarakit*

*Study funded by the National Health and Medical Research Council (APP1033079 and APP1039072) and PhD tenure: Elizabeth and Henry Hamilton-Browne Scholarship, Faculty of Medicine, Sydney Medical School*.

